# Toward a networked central dogma: protein specificity reshapes molecular encounters across evolution

**DOI:** 10.64898/2026.04.21.720012

**Authors:** Zhenzhe Tong, Fei Chen

## Abstract

Since its formulation in 1958, the Central Dogma has provided the organizing framework for molecular biology, describing the informational relationships among DNA, RNA, and protein. Yet, the framework ends at protein synthesis: what proteins do after they are made, how their catalytic specificities determine which molecules actually meet inside the cell, lies beyond its scope. Using a zero-parameter physical model applied to three phylogenetically distant species, we show that physical encounter frequency and metabolic flux are decoupled: raw collision rates predict reaction importance only weakly and show no consistent link to gene essentiality. Enzymatic selectivity bridges this gap through three distinct, species-conserved rescue patterns that persist across more than a billion years of evolution. The cell’s most critical metabolic hubs are, paradoxically, its physically least conspicuous ones— rescued from the collision background by enzymatic precision alone. We term this downstream, measurable layer the Networked Central Dogma: a complement to the classical schema that connects gene expression to the physical organization of cellular chemistry.

## Introduction

The Central Dogma, first articulated by Crick in 1958 and formalized in 1970, established the directional flow of genetic information from DNA to RNA to protein [1, 2]. For over half a century this linear schema has served as the organizing principle of molecular biology. Yet the post-genomic era has revealed rich layers of regulation that the original schema was not intended to cover: transcription factors feed back onto DNA, non-coding RNAs modulate translation at multiple levels, and non-genetic molecules such as metabolites, cofactors and lipids exert pervasive influence on information fidelity and flux through allosteric modulation, covalent modification and phase separation [3–5]. These advances show that the Central Dogma, while precise about template-directed transfer, leaves a downstream question open: once proteins are produced, how do they alter which molecular encounters are favored and which are suppressed?

Living cells contain components whose physical-chemical properties (concentration, diffusivity, half-life) span orders of magnitude, yet network biology traditionally treats all nodes as abstract entities distinguished only by connectivity [6–8]. This topology-only view cannot explain why molecules sharing the same compartment differ by orders of magnitude in encounter frequency. Existing frameworks each capture part of this complexity (Boolean logic, ODE kinetics, scale-free topology, constraint-based flux analysis; Supplementary Table S5) [9–12], but none simultaneously offers zero fitted parameters, explicit molecular physics and full interpretability. The gap is not merely technical: without a parameter-free physical baseline, it is impossible to distinguish the contribution of molecular physics from the contribution of evolved enzymatic specificity.

Our starting point is that no reaction can proceed unless its participants first meet in the right place. The Central Dogma specifies which proteins are made and in what quantity, but says nothing about the probability that a given enzyme will encounter its substrate in a crowded, compartmentalized cell. We address this gap with a zero-parameter three-layer model, Ψ × *J* × *S* (Fig. 1): compartmental gating (Ψ), Smolu-chowski collision frequency (*J*), and enzymatic selectivity (*S*). All inputs are drawn from independently measured data; no parameter is adjusted to fit biological outcomes, so the model can serve as a genuine physical baseline against which the contribution of evolved enzymatic specificity can be directly measured. We apply the model to three metabolites (GTP, acetyl-CoA, ATP) selected for their high biological centrality (each participates in >100 KEGG reactions across all three species) and contrasting physical encounter profiles (Ψ × *J* values spanning two orders of magnitude). Applied across three species spanning prokaryotes and eukaryotes (*E. coli, S. cerevisiae*, mammalian cells), the analysis yields four observations: (i) physical encounter frequency is uncorrelated with metabolic flux, exposing a gap that pure physics cannot close; (ii) each hub metabolite exhibits a distinct enzymatic rescue signature (nucleotide-specific for GTP, redox-module-wide for acetyl-CoA, precision-local for ATP) that mirrors its known biological identity; (iii) citrate synthase emerges as the convergence point of all three encounter axes; and (iv) the prokaryote-to-eukaryote transition transforms a flat encounter space into a modular architecture.

**Figure 1:**
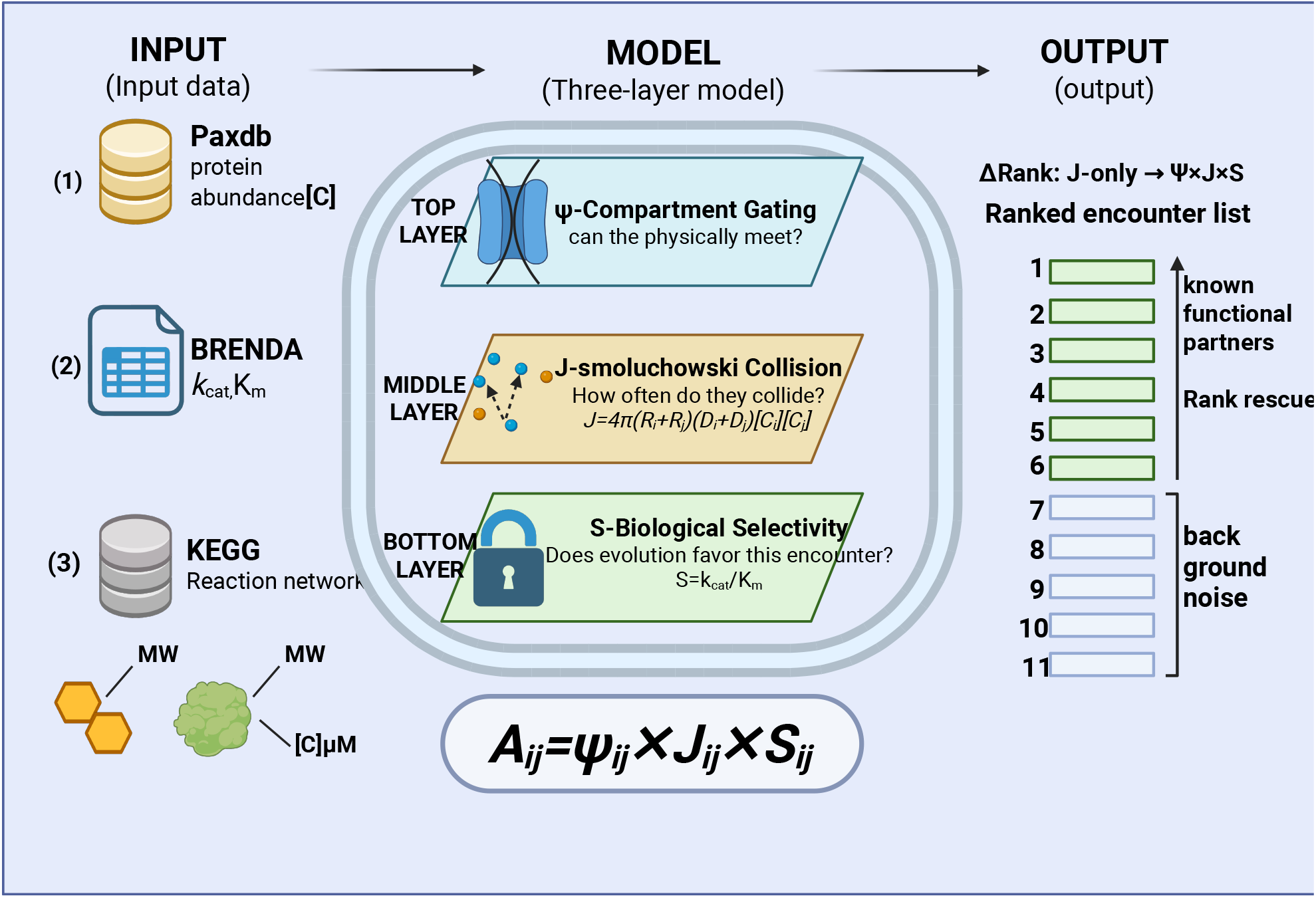
The Ψ × *J* × *S* three-layer collision model. Four independently measured inputs (protein abundance from PaxDb, metabolite concentration from quanti-tative metabolomics, enzyme kinetics from BRENDA, and compartment topology from UniProt/KEGG) yield encounter propensity for every protein–metabolite pair with zero fitted parameters. Ψ: compartmental gating (can they physically meet?); *J* : Smolu-chowski collision frequency (how often do they collide?); *S*: biological selectivity (does evolution favor this encounter?). The rank shift between *J* -only and full-model scoring quantifies how much enzymatic specificity re-ranks each encounter relative to the physics-only baseline.

These observations converge on a single conclusion. Physical encounter frequency and metabolic importance are decoupled, yet enzymatic selectivity bridges this gap in a manner that is conserved across species separated by more than a billion years of evolution. The protein layer, in other words, does not merely execute the instructions encoded in DNA: it actively reorders the physical encounter space of the cell. We term this coupling the “Networked Central Dogma”, a downstream, independently measurable complement to the classical schema that links gene expression to the physical organization of cellular chemistry.

## Results

### Collision frequencies encode partial functional information

The Smoluchowski collision frequency *J*_*ij*_ (Eq. 3) provides a physics-only baseline for molecular encounter propensity. Across three species, *J*_*ij*_ spans approximately eight orders of magnitude, dominated by concentration: ATP, glutamate and other abundant metabolites form the densest encounter neighborhood, whereas low-abundance enzymes and large-molecular-weight complexes occupy the sparse periphery. Ranking all pairs by *J*_*ij*_ alone achieves AUC0.70–0.76 (*n*_pos_ = 509–609; Supplementary Table S3), but drops to 0.43–0.69 for protein–metabolite pairs (*n*_pos_ = 10–11). The full model (Ψ× *J* × *S*) raises all-pair AUC to 0.74–0.80; with matched kinetics only (*S*_kineticsOnly_), protein–metabolite AUC reaches 0.91–0.95 (Fig. 2b). This jump reflects a cliff-like *S*-value separation: non-partners receive *S* = 1, while known enzyme–substrate pairs retain *k*_cat_*/K*_m_ (10^4^–10^8^ M^−1^ s^−1^), creating a four-to-eight order-of-magnitude gap between signal and background. Including KEGG co-occurrence (*S*_full_) actually degrades protein–metabolite classification (AUC0.50–0.71) because cross-species fallback dilutes the kinetic signal. The model’s discriminative power therefore comes from precisely matched enzyme kinetics, not from pathway annotations.

**Figure 2:**
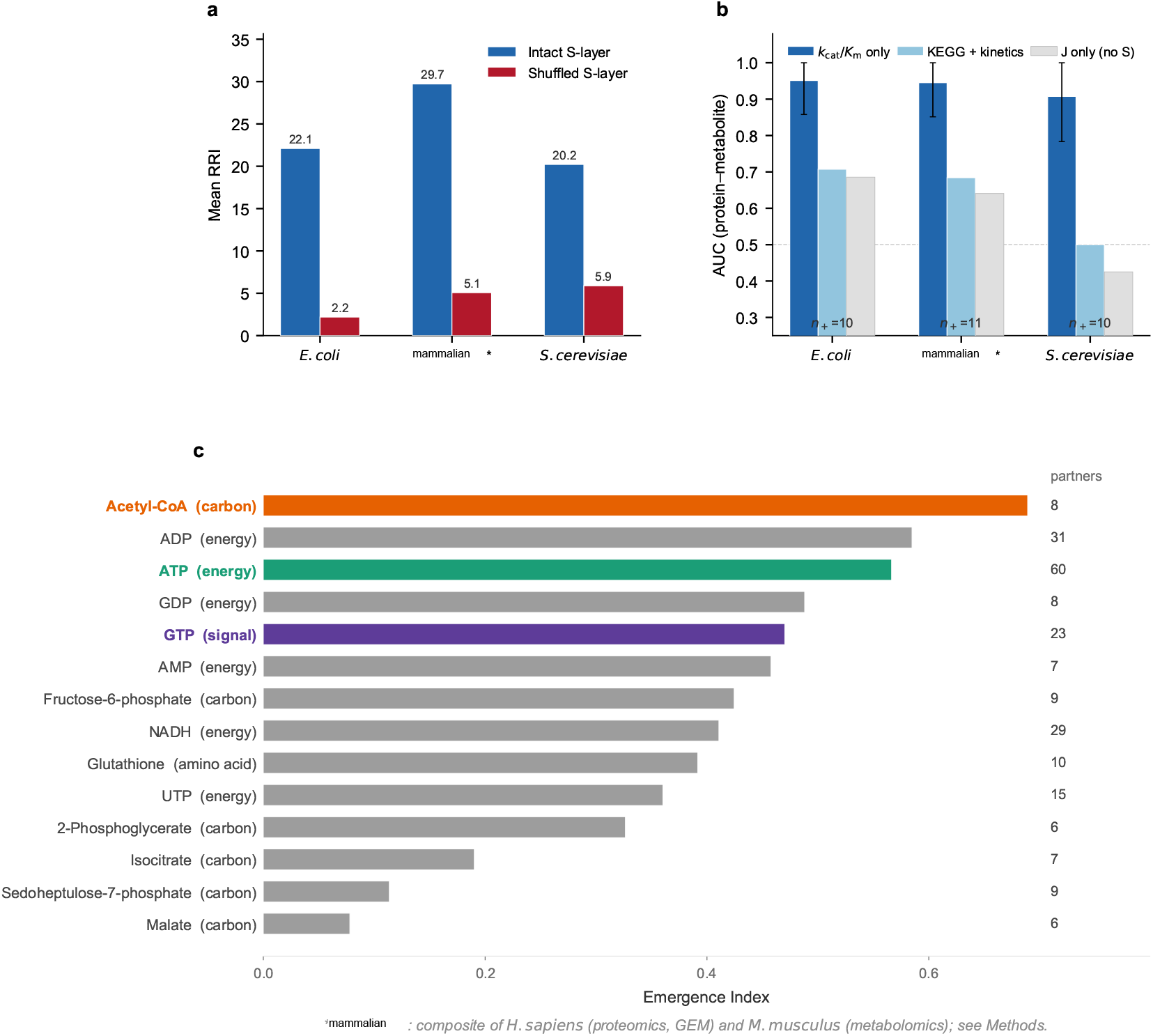
Model validation and star molecule selection. (a) Mean RRI for intact vs. shuffled S-layers (*N* = 3 species, 100 shuffles each); shuffling destroys > 80% of rescue (Wilcoxon *p* < 10^−37^). (b) AUC for classifying known protein–metabolite pairs under three S-layer variants: *S*_kineticsOnly_ (AUC0.91–0.95), *S*_full_ (AUC0.50–0.71), *J* -only (AUC0.43–0.69). Error bars: Hanley–McNeil 95% CI; *n*_+_ = 10–11 (kinetics) or 509–609 (all-pairs). (c) Emergence Index ranking (*N* = 14 metabolites shown); GTP, acetyl-CoA and ATP top three orthogonal axes. Numbers beside bars: validated enzyme partners. *mammalian: composite (see Methods).

### Physical encounter frequency decouples from metabolic flux

If collision frequency alone determined metabolic activity, high-*J*_*ij*_ pairs should also carry high metabolic flux. To test this, we computed the Spearman rank correlation between log_10_ *J*_*ij*_ and log_10_ |FBA flux| for all reaction-associated molecular pairs with non-zero flux in each species’ genome-scale metabolic model (GEM). The correlation was negligible in all three species: *E. coli ρ* = +0.10 (*n* = 540, *p* = 0.021), mammalian *ρ* = −0.14 (*n* = 418, *p* = 0.004), and yeast *ρ* = −0.01 (*n* = 388, *p* = 0.867). Colliding often does not mean reacting; a molecule pair can rank high in encounter frequency yet carry negligible flux, or the reverse. The S-layer (*k*_cat_*/K*_m_) redirects encounter priorities toward functional partnerships.

### Three star molecules under the collision lens

We define the Rank Rescue Index (RRI) as RRI_*ij*_ = rank_*J*_ (*i, j*)−rank_3*L*_(*i, j*): the rank gain from collision-only to full three-layer scoring. GTP, acetyl-CoA and ATP rank highest in a composite Emergence Index (Fig. 2c) and span three orthogonal metabolic axes (Table 1; Fig. 3).

**Table 1:** Three star molecules examined under the collision lens. Values are cross-species means ± SE (*n* = 3: *E. coli, S. cerevisiae*, mammalian). All species-level Wilcoxon *p* < 10^−37^ for S-shuffle Δ.

|  | GTP | Acetyl-CoA | ATP |
| --- | --- | --- | --- |
| Role | Signal relay | Carbon router | Energy currency |
| RRI ± SE | $12.50 \pm 3.77$ | $41.61 \pm 4.74$ | $17.92 \pm 2.54$ |
| S-shuffle $\Delta$ | $10.88 \pm 4.21$ | $32.11 \pm 5.97$ | $15.90 \pm 2.43$ |
| Concordance | 92.7% | 92.1% | 86.0% |
| $\sigma_{\text{break}}^{\text{a}}$ | $\approx 1.0$ | $> 5.0$ | $\approx 5.0$ |
| Specificity | $p \leq 0.01^{\text{b}}$ | $p \leq 0.015^{\text{c}}$ | $\Delta\text{rank} = 24\text{--}41^{\text{d}}$ |

<sup>a</sup> Min. $\sigma$ where any species exceeds 5% rank flip. GTP: *S. cerevisiae* most vulnerable.
<sup>b</sup> Guanine-nucleotide specificity bootstrap.
<sup>c</sup> Redox-module permutation test.
<sup>d</sup> Boundary stress test.
Mammalian: composite of *H. sapiens* (proteomics, GEM) and *M. musculus* (metabolomics); see Methods.

**Figure 3:**
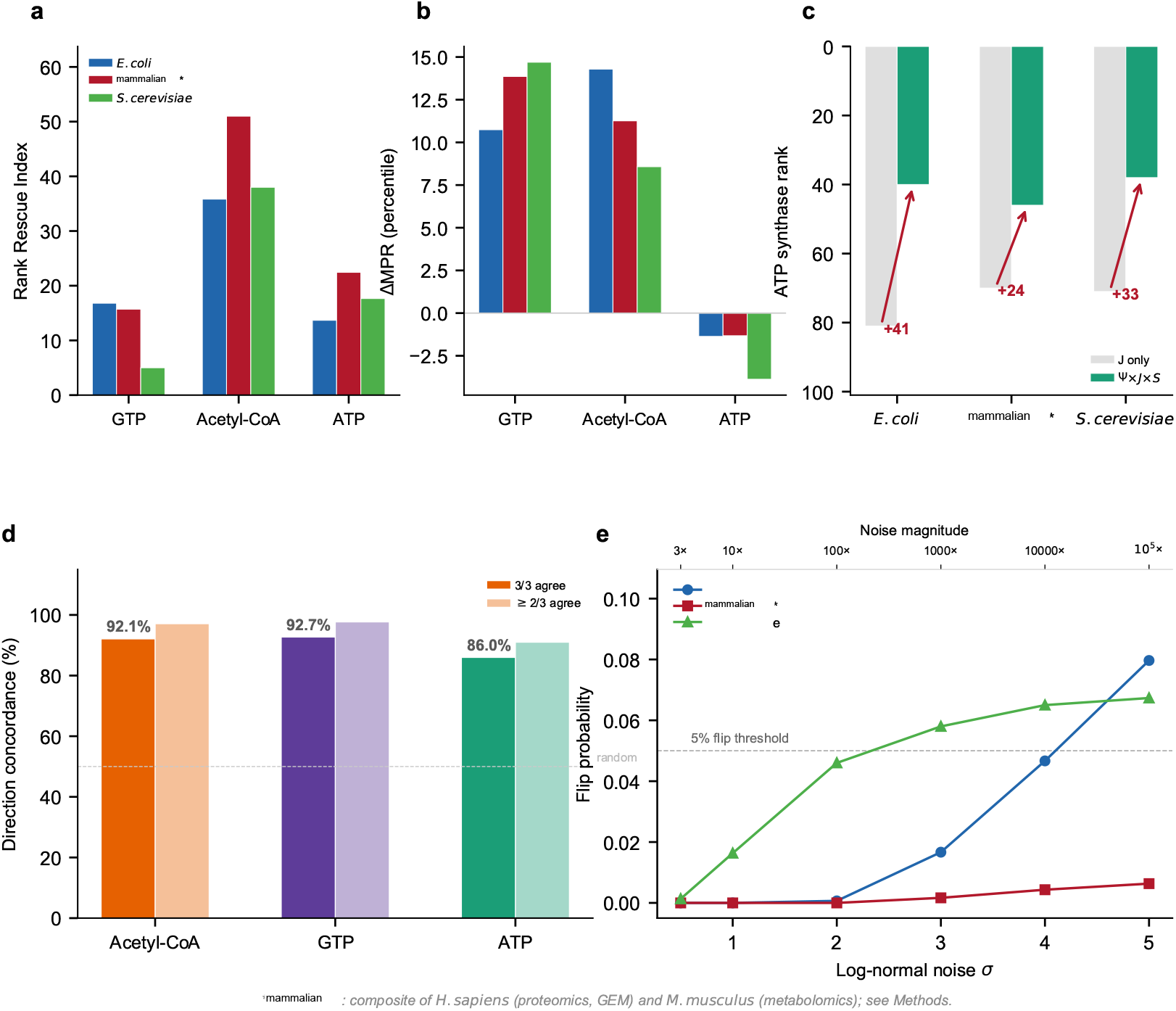
Star molecule portraits and cross-species conservation. (a) RRI per star molecule and species (*N* = 3); acetyl-CoA is the most hidden yet most rescued (RRI = 41.6 ± 4.7 SE), reflecting its paradoxical low physical encounter frequency despite high functional centrality. (b) Global percentile shift (ΔMPR); ATP shows a negative shift (the “hub ranking curse”: molecules already dominating collision space cannot gain further rank). (c) ATP synthase (*α*-subunit) extracted despite hub saturation (Δrank = +24 to +41; *N* = 3 species). FBA gene deletion of *atpA* reduces growth to 32.5% of wild type, confirming functional significance. (d) Direction concordance across species (*n* = 3,422– 3,660 shared metabolite-pair anchors); dashed line = 50% random expectation. Same-sign fraction ranges from 86% (ATP) to 93% (GTP), consistent with conserved encounter architecture across three distant species. (e) Log-normal noise injection (*N* = 1,000 Monte Carlo replicates per *σ*); all species remain below the 5% flip threshold up to *σ* ≈ 3 (∼1,000 × perturbation), far exceeding typical measurement uncertainty. *mammalian: composite of *H. sapiens* and *M. musculus* data (see Methods).

#### GTP: the physically hidden signal relay

In the collision-only ranking, GTP is unremarkable: its concentration and diffusion coefficient place it mid-range among nucleotides. Yet the S-layer rescues GTP with a specificity that distinguishes it from every other NTP: a bootstrap test against ATP, CTP and UTP confirms that GTP’s rank rescue is locked to the guanine-nucleotide pool (Wilcoxon *p* ≤ 0.01 in all three species), excluding a generic nucleotide effect. This fits a molecule whose biological roles (G-protein signaling, GTPase regulation, translational elongation [13]) all require partner selectivity that diffusion physics alone cannot provide.

#### Acetyl-CoA: the most hidden, most rescued metabolite

Acetyl-CoA occupies a paradoxical position: it is physically concealed (median concentration 2–4× lower than glycolysis intermediates across species; Supplementary Table S2), yet it shows the highest enzymatic rescue among all metabolites (RRI = 41.61 ± 4.74 SE) and strong cross-species concordance (92.1 %). Its rescue does not act on individual partners alone: a Wilcoxon signed-rank test shows that the entire redox cofactor module (NADH, NADPH, NADP^+^, FAD) is co-elevated (*p ≤* 0.015 in all species), a pattern that parallels acetyl-CoA’s position at the junction of glycolysis, the tricarboxylic acid (TCA) cycle and fatty acid synthesis [14, 15]. In other words, the carbon hub with the highest functional centrality also has the lowest physical encounter frequency; the S-layer compensates with the largest rank rescue observed for any metabolite.

#### ATP: the hub ranking curse and precision extraction

ATP presents the opposite challenge: it physically dominates the encounter space (26–65 known partners across species, crowding ratio 0.74–0.78), leaving no room for global rank improvement (ΔMPR = −1.3 to −3.9; Fig. 3b). This “hub ranking curse” is mathematically inevitable when a molecule already occupies the densest region of collision space. Yet within this noise, one signal survives: ATP synthase, among the most conserved enzyme complexes in biology [16, 17], is elevated by 24–41 rank positions in every species (Fig. 3c). FBA gene deletion confirms this is not a ranking artifact: *atpA* deletion reduces growth to 32.5% of wild type. Even when a molecule saturates encounter space, the model can still pick out individual partnerships whose specificity signal is strong enough.

#### Citrate synthase: the triple crossroads

Supplementary Table S9 lists the specific molecular partners whose encounter priority is most affected by enzyme specificity. One result stands out: citrate synthase, the TCA cycle entry enzyme, is independently rescued by all three star molecules in all three species (Δrank = +46 to +77). This convergence is not coincidental: citrate synthase sits at the metabolic node where signal relay (GTP, via succinyl-CoA synthetase coupling), carbon flux (acetyl-CoA, as direct substrate) and energy transduction (ATP, via TCA-driven oxidative phosphorylation) all intersect [18, 19]. Single-gene deletion of *gltA* in iML1515 yields zero growth, confirming that encounter-space convergence tracks functional indispensability.

#### Bidirectional validation: proteins as anchors

All analyses above use metabolites as anchors and rank proteins by encounter propensity. To test whether the restructuring operates symmetrically, we repeated the atlas using 240 proteins as anchors (80 per species) and ranking metabolites. In *E. coli* and yeast, protein-anchored RRI is positive (+1.61 and +1.42, respectively), confirming that the S-layer elevates known substrates from both ends of the protein–metabolite interface. Key enzymes, including cytochrome *c* oxidase, RNA polymerase and citrate synthase, show the strongest enrichment of known substrates in their top-10 encounter partners (permutation *p* < 0.001; hits@10 ratio = 1.0), independently recapitulating the star-molecule findings from the metabolite perspective. Mammalian protein-anchored RRI is slightly negative (−0.84), attributable to 45% of mammalian proteins having zero known metabolite partners in current databases (Discussion).

#### Robustness

Randomly reassigning enzyme–substrate pairings in the S-layer drops mean RRI from 20–30 to 2–6 (*p* < 10^−37^; Fig. 2a), showing that the rescue signal is not a statistical artifact of network topology. The model holds under leave-one-out removal of individual species, a tenfold parameter scan, and log-normal input noise (Supplementary Tables S4–S7; Fig. 3e): predictions degrade only when input perturbations exceed *σ* ≈ 3, roughly 1,000× the typical measurement error.

#### Gene essentiality enrichment

FBA single-gene deletion (independent of the HTS framework) confirms that star-molecule genes are enriched for essentiality: ATP-associated genes in *E. coli* (16.8% vs. 11.8%; OR = 1.52, *p* = 0.010) and GTP-associated genes in yeast (42.9% vs. 11.7%; OR = 5.68, *p* = 0.004). Genes at the intersection of two star-molecule networks are essential at ∼36% (∼3× background), with a dose–response gradient (*ρ* = +0.063, *p* = 0.014). Recon3D was excluded due to its anomalously high FBA growth rate.

### Star-molecule encounter patterns are consistent across phylogenetically distant species

Patterns observed in a single species could be idiosyncratic. We therefore asked whether the encounter portraits of GTP, acetyl-CoA and ATP hold across evolutionarily distant organisms whose parameters were measured independently.

#### Global conservation of rescue direction

For 3,422–3,660 shared metabolite-pair anchors across species, we computed Spearman rank correlations of pairwise RRI values: *E. coli* vs. mammalian *ρ* = 0.70 (*p* < 10^−300^, *n* = 3,660), *E. coli* vs. yeast *ρ* = 0.67 (*p* < 10^−300^, *n* = 3,660), and mammalian vs. yeast *ρ* = 0.84 (*p* < 10^−300^, *n* = 3,422). The same-sign fraction ranges from 93.0% to 95.3%, far exceeding the 50% expected by chance (Fig. 3d). Per-hub concordance is highest for GTP (92.7%) and lowest for ATP (86.0%), mirroring the gradient observed in the rescue analysis above: the metabolite whose rescue is most structurally constrained (signal routing) is also the most conserved, while the physically saturated hub (energy currency) shows the most inter-species variation. Even the collision-only layer (*J*) is partly conserved: *ρ* = 0.50–0.52 between prokaryote and eukaryote, rising to *ρ* = 0.80 within eukaryotes [20, 21]; yet the full three-layer model consistently adds Δ*ρ* = +0.07 to +0.17, suggesting an additional layer of conserved biological organization.

#### Physical concealment versus functional necessity

The data reveal an asymmetry between metabolite classes. Redox cofactors (NAD^+^, NADH, NADP^+^, NADPH, FAD) have measurably lower intracellular concentrations than glycolysis intermediates (median 65–183 *µ*M vs. 375–1 540 *µ*M across species; Supplementary Table S2), making them physically hidden in the encounter ranking. Yet these are precisely the molecules that serve as electron-transfer “wires” connecting metabolic modules. Accordingly, redox-cofactor pairs show consistently positive mean RRI, whereas glycolysis and TCA cycle intermediates, physically prominent and already occupying the dense core of collision space, show negative mean RRI. This contrast is conserved across all three species.

Abundant intermediates need no rescue because collision physics already places them in the encounter-rich zone; cofactors, by contrast, must bridge concentration regimes to link metabolic subsystems. Their concealment creates a “rescue demand” that enzyme specificity fills. This asymmetry suggests that enzymatic rescue is distributed according to the gap between a molecule’s physical encounter frequency and its metabolic importance.

### From flat encounters to modular architecture: an evolutionary trajectory

Newman modularity *Q* [22] (top 5% of *A*_*ij*_, vs. 1 000 degree-preserving random graphs) reveals a sharp prokaryote–eukaryote divide: *E. coli Q* = −0.006 (*z* = 1.40, *p* = 0.096; flat, concentration-dominated); yeast *Q* = 0.034 (*z* = 9.37, *p* < 10^−4^); mammalian *Q* = 0.047 (*z* = 10.03, *p* < 10^−4^).

This transition parallels organelle acquisition [23, 24] (Ψ-layer partitioning) and enzyme-family expansion (*S*-layer enrichment), turning a flat encounter space into a modular one. Protein-encoded specificity thus progressively restructures the molecular encounter space as cellular complexity increases.

## Discussion

The collision model provides a single physical framework for three well-known hub metabolites. GTP’s nucleotide-specific rescue matches its signaling selectivity [13]; acetyl-CoA’s paradox (lowest physical encounter, highest rescue) fits its junctional role between glycolysis, TCA and fatty-acid synthesis; and the 86–93% cross-species concordance places these encounter junctions among the most conserved features of cellular organization. Two factors explain this concordance: (i) absolute metabolite concentrations are conserved across kingdoms [21, 25], and (ii) core-pathway enzymes face stronger selection pressure (*k*_cat_*/K*_m_ *∼* 30× higher; [26]). For ATP, the hub-ranking effect suppresses global rescue, yet ATP synthase [27, 28] still emerges, and a hub-removal stress test confirms that deleting energy-currency nodes collapses hits@10 by > 60%.

Independent evidence corroborates each finding. The collision-only ranking is itself partly conserved (*J* -rank *ρ* = 0.50–0.52 cross-domain, 0.80 within eukaryotes), consistent with conserved metabolite concentrations [25]. *k*_cat_*/K*_m_ tracks flux requirements rather than substrate abundance [26, 29], explaining the encounter–flux decoupling (|*ρ*| *≤* 0.14). FBA shadow prices (Supplementary Table S8) show the GTP/ATP ratio rising from prokaryote to eukaryote (1.0 → 1.12 → 2.00), tracking the encounter-space hierarchy. Gene essentiality enrichment [32] shows a dose–response gradient (∼ 3× background at the intersection of two hub networks), though the bottleneck molecule differs by species (ATP in *E. coli*, GTP in yeast), suggesting that enzymatic specificity operates as a conserved evolutionary constraint [33] even as its focal target shifts.

Taken together, these patterns reveal a layer of cellular organization that the Central Dogma does not address. Physical encounter frequency sets a baseline that evolution has systematically overridden: the protein layer reorders encounter space through kinetic specificities that are themselves genetically encoded. We term this coupling the “Networked Central Dogma” (Fig. 4). Three lines of quantitative evidence support this view: rescue direction is conserved across three distant species; rescue is concentrated on physically concealed cofactors rather than abundant ones; and the prokaryote-to-eukaryote transition marks a shift from concentration-dominated to specificity-driven encounter organization.

**Figure 4:**
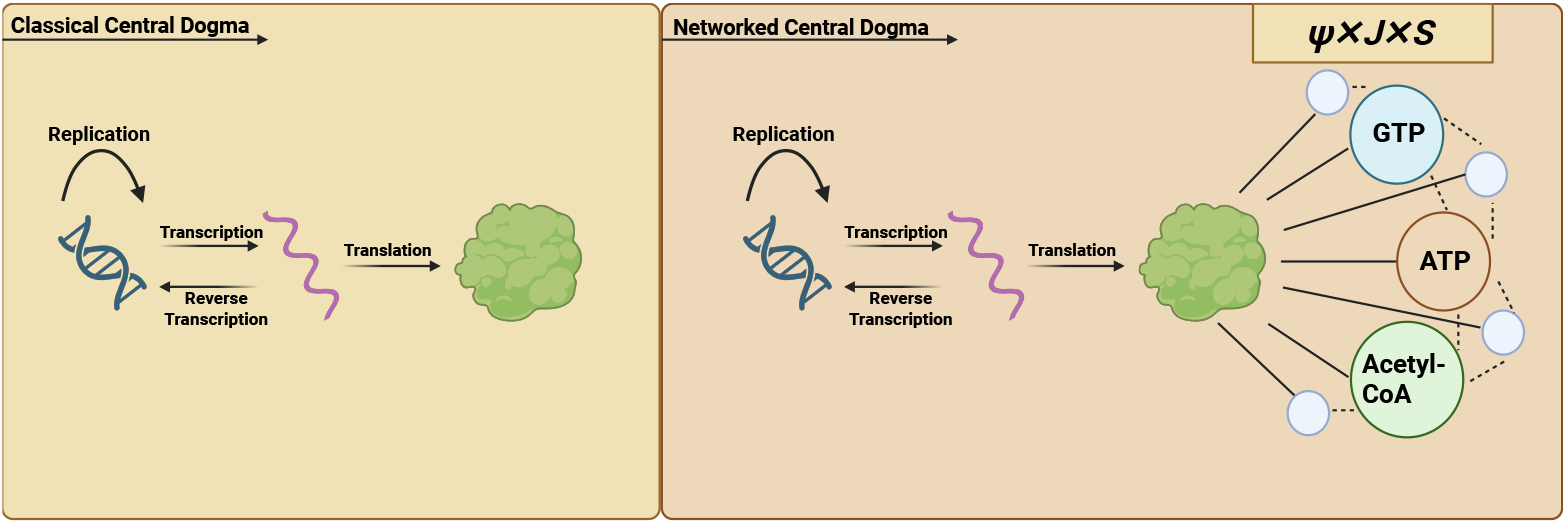
The Networked Central Dogma. Left: The Classical Central Dogma (DNA → RNA → Protein) describes template-directed information flow but is silent on downstream molecular encounters. Right: The collision lens (Ψ × *J* × *S*) reveals a previously unmeasured layer: proteins reorder the physical encounter space through their catalytic specificities (*k*_cat_*/K*_m_). Three hub molecules (GTP, ATP, Acetyl-CoA) exemplify three distinct modes of encounter restructuring: nucleotide-specific rescue (GTP), redox-module co-elevation (Acetyl-CoA), and precision extraction from a saturated background (ATP). Rescue directions are consistent across all three species (86–93% concordance), coinciding with a prokaryote-to-eukaryote shift from flat encounter spaces (*Q* ≈ 0) to modular architectures (*z* > 9, *p* < 10^−4^). Thick lines = high-specificity (enzymatically favored) encounters; thin lines = low-specificity (collision-dominated) encounters. All quantitative values are from Figs. 2–3 and Table 1.

The implications extend beyond the three focal metabolites. If the protein layer systematically reshapes encounter space, then the physical-chemical niche of each catalytic strategy should itself be a conserved feature of life. The EC-class analysis confirms this prediction: each enzyme class occupies a distinct and reproducible region of the saturation–gap parameter space across ten species (Supplementary Table S10; Cliff’s δ = 0.36, *p* = 7.8 × 10^−10^). For metabolic engineering, the concealment–necessity asymmetry predicts that synthetic pathways requiring non-native cofactor wiring will incur higher metabolic burden, a testable hypothesis that connects encounter physics directly to design constraints.

The prokaryote-to-eukaryote transition (*E. coli Q* ≈ 0; eukaryotes *Q* > 0, *z* > 9) reflects two innovations: organelle acquisition (partitioning Ψ) and enzyme-family expansion (enriching *S*), shifting encounter space from concentration-dominated to specificity-driven. This complements topological explanations of metabolic modularity [34–36] by grounding it in physical parameters.

Three principal caveats apply. First, the model is static: it captures time-averaged encounter propensity but not temporal dynamics, allosteric regulation, or ternary complexes; when matched kinetic data are unavailable, the S-layer falls back on KEGG co-occurrence, which dilutes classification power (Supplementary Table S3). Second, the mammalian species is a human–mouse composite (see Methods); a fully species-matched dataset remains desirable, though substitution sensitivity analysis shows negligible impact (*ρ* = 0.97; Supplementary Fig. S5). Third, a direct closure test from encounter priority to gene essentiality was not confirmed (*E. coli p* = 0.013, direction opposite to expectation), consistent with the Drummond–Wilke effect [37, 38]: the encounter model captures pair-level proximity, not gene-level dispensability.

Time-resolved proteomics could extend the static encounter map into a dynamic picture. Integration with genome-scale models would couple encounter propensity with flux feasibility. The concealment–necessity asymmetry predicts that synthetic pathways requiring non-native cofactor wiring should incur higher metabolic burden; testing this prediction would supply direct experimental evidence for the Networked Central Dogma.

## Methods

### Data sources

The master dataset (V41_Master.csv) integrates four public databases. Protein abundances were obtained from PaxDb v4.2 [39] for *E. coli* K-12 (integrated dataset), *S. cerevisiae* S288C, and *Homo sapiens* (whole-organism integrated dataset); ppm values were converted to intracellular *µ*M concentrations using species-specific cell volumes [40] (*V*_E. coli_ = 0.7 fL; *V*_yeast_ = 42 fL; *V*_mammalian_ = 2,000 fL) and mean protein molecular weight from UniProt. Metabolite concentrations were assembled from Park *et al*. (2016) [21] for iBMK (*Mus musculus*, immortalised baby mouse kidney epithelial cells) and supplemented with nucleotide values from Traut (1994) [41]; for *E. coli* and yeast, values from Bennett *et al*. (2009) [25] were used. All concentrations reported in mM were converted to *µ*M (×1 000). Enzyme kinetics (*k*_cat_, *K*_m_, *k*_cat_*/K*_m_) were retrieved from BRENDA 2026.1 [42] and cross-checked with SABIO-RK [43], using EC-number matching; organism-specific entries were preferred, with cross-species proxies used only when no same-organism data were available (all proxied entries labeled in the Data_Source column). For the mammalian species, 34.5% of kinetic entries were organism-matched (*H. sapiens*); the remainder were drawn from cross-mammalian sources (*M. musculus, R. norvegicus*). Genome-scale metabolic models (GEMs) for flux-balance analysis (FBA) were iML1515 for *E. coli* [44], iMM904 for *S. cerevisiae* [45], and Recon3D for *H. sapiens* [46].

#### Mammalian composite dataset

The mammalian species in this study is a composite: protein abundances and the genome-scale model (Recon3D) are from *H. sapiens*, while metabolite concentrations are from *M. musculus* iBMK cells [21]. Three lines of evidence justify this combination: (i) human–mouse gene-level conservation exceeds 95% for core metabolic enzymes [47]; (ii) the physical inputs to the model (molecular weight, diffusion coefficient, compartment assignment) are identical for orthologous proteins; and (iii) mixing two mammalian sources introduces a noise floor that makes any observed signal a conservative lower bound. A substitution sensitivity analysis (Supplementary Fig. S5) confirms that replacing mouse metabolite concentrations with human estimates changes fewer than 3% of top-ranked encounters (*ρ* = 0.97 between composite and human-only rankings).

### Three-layer model

#### Ψ-layer (compartmental gating)

The encounter propensity between protein *i* and metabolite *j* is defined as a product of three independently computed layers:

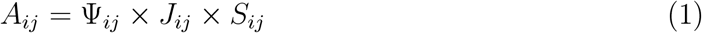

If two molecules share the same normalised compartment label, Ψ_*ij*_ = 1. For cross-compartment pairs, a sigmoidal penalty is applied:

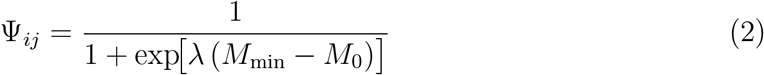

where *M*_min_ = min(*M*_*w,i*_, *M*_*w,j*_), *M*_0_ = 40 000 Da is the midpoint and *λ* = 0.001 Da^−1^ is the slope. This formulation assigns higher gating to small molecules (which diffuse between compartments more readily) and progressively penalises large-protein cross-compartment encounters. Compartment annotations were obtained from UniProt (proteins) and KEGG (metabolites), normalised to five canonical labels (Cytoplasm, Membrane, Mitochondria, Nucleus, Other).

#### *J* -layer (Smoluchowski collision frequency)

The bimolecular diffusion-limited encounter rate is computed from the Smoluchowski equation [48] for two spheres in solution:

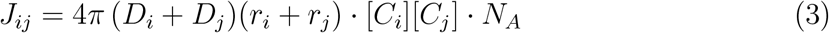

where *D*_*i*_ is the translational diffusion coefficient estimated from the Stokes–Einstein relation *D* = *k*_*B*_*T*/(6*πηr*), *r*_*i*_ is the hydrodynamic radius estimated from molecular weight using sphere-volume scaling 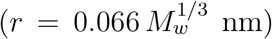, [*C*_*i*_] is the intracellular concentration in *µ*M, and *N*_*A*_ is Avogadro’s number. Cytoplasmic viscosity was set to *η* = *η*_water_(37 °C) × 3 = 0.692 mPa · s × 3 ≈ 2.08 mPa · s, where the crowding factor of 3 is consistent with fluorescence correlation spectroscopy measurements in bacterial and mammalian cytoplasm [49–51]. Temperature was fixed at *T* = 310.15 K (37 °C) for all species. All parameters are independently measured; no fitting is performed.

#### *S*-layer (biological selectivity)

The selectivity score *S*_*ij*_ is assigned hierarchically: (i) organism-matched *k*_cat_*/K*_m_ from BRENDA (preferred); (ii) cross-species proxy *k*_cat_*/K*_m_ when same-organism data are unavailable; (iii) EC-level median *k*_cat_*/K*_m_ when no substrate-specific data exist; (iv) a KEGG [52] reaction co-occurrence score (10^*n*_rxn_^, capped at 10^6^) when no kinetic data are available; (v) a neutral fallback *S* = 1 (no information, reducing the three-layer score to Ψ × *J*). For metabolite–metabolite pairs, KEGG pathway co-occurrence is used; protein–protein pairs receive *S* = 1. Each pair’s *S* source is recorded in the output for traceability.

### Validation experiments

#### S-layer shuffle (HPC-7)

To test causality, the S-layer values were randomly permuted across all molecule pairs while keeping Ψ and *J* intact. 100 independent shuffles × 3 anchor molecules were performed per species using high-performance computing (HPC), and the mean RRI across shuffles was compared to the intact model using a two-sided Wilcoxon signed-rank test.

#### De-leakage (S-layer variant) experiment

Three S-layer variants were constructed to disentangle the contribution of enzyme kinetics from KEGG co-occurrence: *S*_full_ (all information), *S*_noKEGG_ (kinetics + concentration, no pathway labels), and *S*_kineticsOnlyStrict_ (matched *k*_cat_*/K*_m_ values only). Classification performance (AUC) was evaluated on validated protein–metabolite pairs and on all pairs, with 5-fold cross-validation to estimate variance.

#### Log-normal noise injection (HPC-8)

All three input layers were independently perturbed by multiplicative log-normal noise exp(𝒩 (0, *σ*^2^)) for *σ* ∈ {0.5, 1.0, 2.0, 3.0, 4.0, 5.0}, corresponding to ∼3× to ∼10^5^× perturbation. 1 000 replicates per *σ* level were run on HPC, and the flip probability (fraction of replicates where the top-10 ranked metabolites changed relative to the unperturbed model) was recorded. The top-10 ranking remained stable up to *σ* ≈ 3.0 for *E. coli* and mammalian cells and *σ* ≈ 1.0 for yeast (Supplementary Fig. S2).

#### Leave-one-out sensitivity (HPC-6)

Each enzyme and each metabolite were individually removed from the dataset, and the full three-layer ranking was recomputed. The critical fraction was defined as the percentage of removals that caused any top-10 metabolite to change.

#### Encounter–flux decoupling test

FBA was performed using COBRApy [53] on each GEM with default growth-medium constraints. For each reaction-associated molecular pair with non-zero flux, we computed the Spearman rank correlation between log_10_ *J*_*ij*_ and log_10_ |flux|. For mammalian cells, the anomalously high Recon3D growth rate (∼ 377 h^−1^) was addressed by using normalised metabolic headroom (*H*_norm_).

#### Pathway cofactor rescue permutation test

For five defined metabolic modules (redox cofactors, CoA family, energy carriers, TCA cycle, glycolysis), we computed the mean RRI of all within-module KEGG-validated pairs and compared it against a null distribution generated by randomly sampling equal-sized sets from all KEGG-validated pairs (1 000 permutations per species per module). One-sided *p*-values (fraction of permutation means ≥ observed) were reported for positively rescued modules.

#### Gene essentiality enrichment test

To independently validate the star-molecule portraits, we performed genome-wide single-gene deletion using COBRApy [53] on iML1515 (*E. coli*, 1 516 genes [44, 54]) and iMM904 (*S. cerevisiae*, 905 genes [45, 55]). A gene was classified as essential if its deletion rendered FBA infeasible or reduced growth below 5% of wild type. For each star molecule (GTP, acetyl-CoA, ATP), we identified all GEM reactions involving the metabolite and collected the catalyzing genes. Essentiality enrichment was assessed using Fisher’s exact test (one-sided, alternative = “greater”). A dose–response analysis tested whether genes at the intersection of multiple star-molecule reaction sets showed higher essentiality rates, with Spearman rank correlation used for trend significance. Recon3D was excluded from this analysis because its anomalously high FBA growth rate (*∼* 377 h_−1_) yielded only 1 essential gene out of 2 248, precluding meaningful enrichment testing.

#### Newman modularity

The encounter network was thresholded at the top 5% of *A*_*ij*_ scores and binarised. Newman modularity *Q* was computed and compared against 1 000 degree-preserving random graphs to obtain *z*-scores and empirical *p*-values.

### Statistical analysis

All statistical tests were performed in Python 3.13 using SciPy 1.14 and scikit-learn 1.6. Two-sided Wilcoxon signed-rank tests were used for paired comparisons (intact vs. shuffled rankings) with no correction for ties. AUC was computed using `sklearn.metrics.roc_auc_score;` 95% confidence intervals were estimated using the Hanley–McNeil (1982) formula [56]:

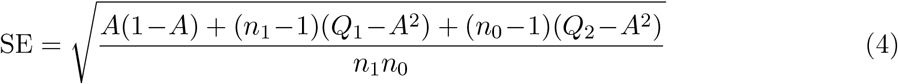

where *Q*_1_ = *A*/(2 *− A*) and *Q*_2_ = 2*A*^2^/(1 + *A*). GTP specificity was assessed by bootstrap comparison against ATP, CTP, and UTP RRI values (10 000 resamples). All random seeds were fixed `(random.seed(42), np.random.seed(42))` for reproducibility. For sample sizes *<* 10, non-parametric tests were used exclusively and effect sizes reported alongside p-values.

### Use of AI tools

Claude (Anthropic) was used for writing assistance and language polishing during manuscript preparation. All scientific content, data analyses, interpretations and conclusions are the sole responsibility of the human authors.

## Data availability

All raw data used in this study are from publicly available databases: protein abundances from PaxDb v4.2 (https://pax-db.org/), metabolite concentrations from published literature [21, 25, 41], enzyme kinetics from BRENDA 2026.1 (https://www.brenda-enzymes.org/) and SABIO-RK (http://sabiork.h-its.org/), and genome-scale metabolic models from BiGG Models (http://bigg.ucsd.edu/). The processed master dataset (V41_Master.csv) and all intermediate data files are available at https://github.com/important-never/HTS-CollisionModel.

## Code availability

All custom Python code used to generate the results reported in this study is available at https://github.com/important-never/HTS-CollisionModel. The repository includes scripts for three-layer model computation, cross-species analysis, validation experiments (S-layer shuffle, noise injection, leave-one-out), and figure generation.

## Acknowledgements

We thank the developers and maintainers of PaxDb, BRENDA, SABIO-RK, KEGG, and UniProt for making their databases publicly available, and the COBRApy team for open-source flux-balance analysis tools. Genome-scale metabolic models were obtained from the BiGG Models repository (iML1515, iMM904, Recon3D).

## Author contributions

F.C. conceived and led the study. Z.T. performed the analyses and wrote the initial draft.

F.C. and Z.T. revised the manuscript and approved the final version.

